# Whole-genome prediction of bacterial pathogenic capacity on novel bacteria using protein language models, with PathogenFinder2

**DOI:** 10.1101/2025.04.12.648497

**Authors:** Alfred Ferrer Florensa, Jose Juan Almagro Armenteros, Rolf Sommer Kaas, Philip Thomas Lanken Conradsen Clausen, Henrik Nielsen, Burkhard Rost, Frank Møller Aarestrup

## Abstract

Infectious diseases continue to be a leading cause of mortality and pose a significant global health threat. Thus the development of tools for surveillance and early detection of emerging pathogens is needed. In this study, we introduce PathogenFinder2, a novel predictor of bacterial pathogenic capacity in humans, available through an online server (http://genepi.food.dtu.dk/pathogenfinder2), or as a standalone program (https://github.com/genomicepidemiology/PathogenFinder2). The model, using protein language models for whole-genome phenotype prediction, surpasses the performance of previous methods, especially for novel bacterial taxa, while being taxonomy-agnostic and alignment-free. At the same time, it predicts the importance of each protein for the pathogenic capacity. This output might aid in characterizing potential pathogens, it readily identifies new candidates for virulence factors and vaccine targets, and offers insights into infection metabolic pathways. Furthermore, we introduce the Bacterial Pathogenic Landscape, revealing distributions related to the host conditions, antagonist bacteria, infection site, or habitat.

## 1 Introduction

Human-bacterial interactions vary significantly. While some bacteria cannot colonize humans, others engage in non-damaging or even beneficial interactions like mutualism and commensalism, or harmful ones as pathogens[11, 18]. New possible beneficial bacteria are today widely explored for health and commercial purposes, including starter cultures, pre- and probiotics, and industrial applications. Moreover, with anticipated climate changes and greater exploration of natural resources, human exposure to novel bacterial taxa are also expected to rise. Therefore, it is crucial to develop tools to distinguish harmful bacteria from harmless ones, enabling early detection and timely interventions, even when faced with novel taxa.

Differentiating between pathogenic and non-pathogenic bacteria has been a longstanding challenge for the scientific community, even since Koch’s postulates[21]. The outcome of bacterial contact with humans varies significantly, as evidenced by the ongoing HIV/AIDS epidemic. Infections by bacteria previously deemed to be non-pathogenic in immunocompromised hosts challenged the binary view of pathogenicity, leading to the concept of “opportunistic pathogens”[3]. However, defining this term remains complex, as infection incidence of any type of pathogen is notably higher in immunocompromised hosts; and even individuals with a healthy immune system can be infected by “opportunistic pathogens”[10, 41]. This lack of consensus underscores the host’s role in the pathogen’s success (and its resulting damage). Factors such as the immune system, infection site, and host microbiome will significantly influence the interaction outcome[41, 12].

However, the pathogenic capacity[13] to cause host damage under appropriate conditions must be encoded in the bacterium’s genetic material[57]. Over many years, significant efforts have identified proteins and genes that define the pathogenic capacity and virulence of a bacterium. The extensive availability of bacterial genome data has significantly broadened our understanding of these virulence factors[35]. However, many of these genes are also present in commensal bacteria[27], impeding their use for prediction of bacterial pathogenic capacity.

Traditionally, predicting bacterial pathogenic capacity has often involved inoculating pathogens in animal models. However, varying host ranges among bacteria limit this method’s effectiveness[21]. Another approach is to infer pathogenic capacity from the phenotype of closely related bacteria. This method’s effectiveness is constrained by database completeness, bias, and identification accuracy. Moreover, pathogenic capacity and taxonomy do not always correlate, as some species have both pathogenic and non-pathogenic strains[43]. In fact, the acquisition of a mobile genetic element or a few point mutations can enable a bacterium to cause damage in a human host[51].

Recent methods to predict pathogenicity have relied on machine learning, falling into two categories: protein- and read-based methods. Protein-based methods typically involve three steps: protein prediction, mapping to a protein family database, and prediction based on the hits identified. Current state-of-the-art solutions, such as PathogenFinder[16], BacPacs[7] and WSPC[42], employ various strategies for database construction and prediction, to predict pathogenic capacity of bacterial isolates and identify key proteins involved in pathogenicity. Limitations of those methods include the need for assembled genomes, the exclusion of non-proteomic genetic material, and the reliance on protein alignment, which is constrained by database completeness and alignment analysis. In contrast, read-based methods, such as PaPrBag[17] and DeePac[8], do not require genome assembly or protein prediction, nor do they necessitate mapping or the creation of a protein family database. Drawbacks for such read-based methods include the inability to explain the mechanisms underlying pathogenicity when modeling bacterial sequences.

Here, we introduce PathogenFinder2, a novel protein-based method for predicting bacterial pathogenic capacity using protein Language Models (pLMs), without relying on traditional alignment or mapping techniques. This method enables whole-genome phenotype prediction through a four-step process (Figure 1(a)). By representing bacteria as the ensemble of their embedded proteins using ProtT5[20], we leverage the extensive knowledge from pLMs, and achieve representations that vary at the strain level. This genomic representation serves as the input for the fourth step, a deep neural network (Figure 1(b)). The model firstly utilizes layers with 1D convolutions that address the variable and relatively large length of the input (e.g., *Escherichia coli* has approximately 4,288 coding genes). As the input is learned through convolution kernels, we also expect the model to learn local relationships (e.g., operons or mobile elements). Secondly, an attention layer confers different importance to each protein, as we assume that not all of them will be equally relevant for the prediction.

**Figure 1:**
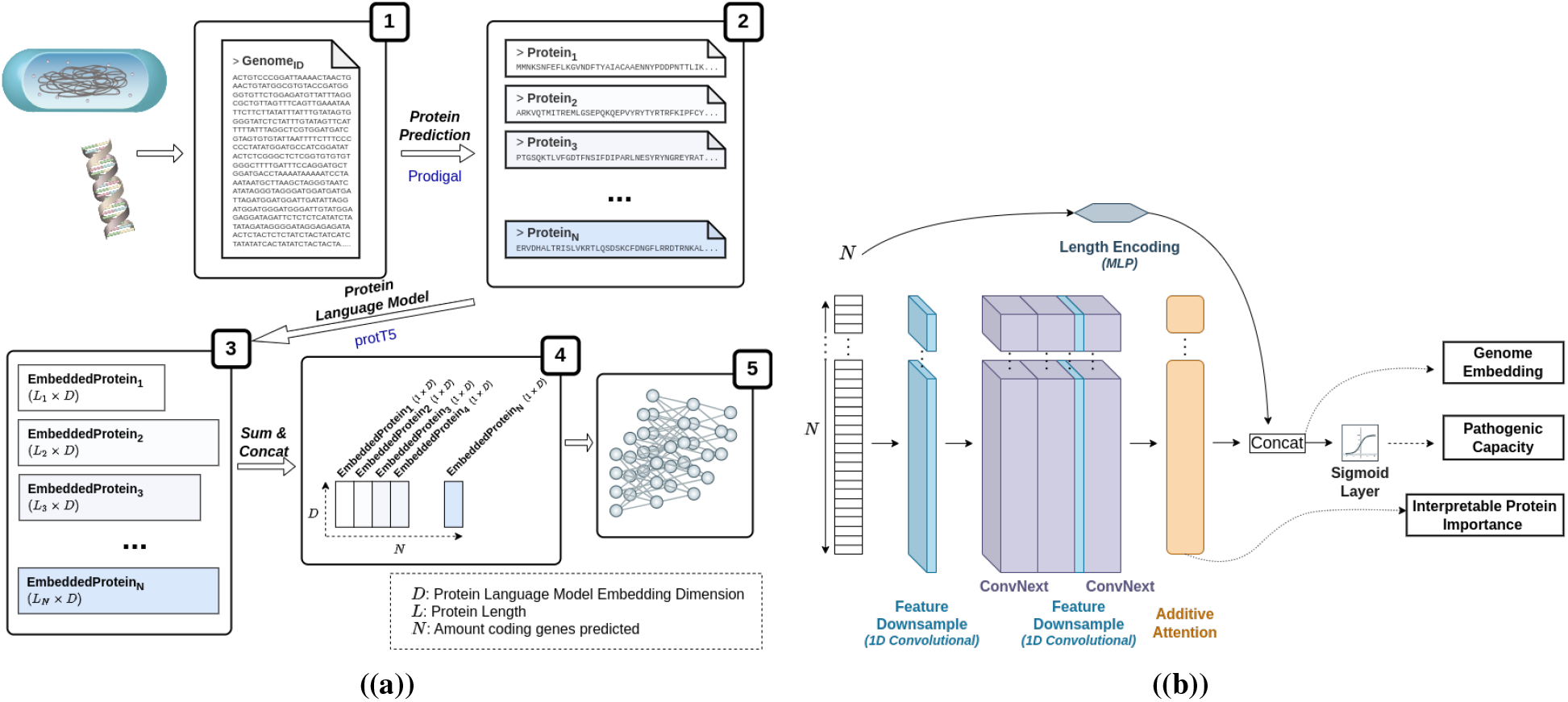
The PathogenFinder2 model. a) Overall pipeline of PathogenFinder2, from the assembled genome until the neural network. The genome is converted into a collection of predicted proteins using Prodigal[28], which are then embedded using the protT5[20] protein language model. The embeddings are summed along the amino acid length axis, resulting in a 1xD vector for each protein (*D* being the dimensions of the protein language model embedding). These vectors are stacked to form a sequence used as input for the deep learning model. b) Schematic of the PathogenFinder2 neural network, comprising two main components: an adaptation of the ConvNext[37] image classifier for sequential data (consisting of 1D Convolutions) and a Bahdanau attention[6] layer to account for unequal protein importance. These components, free from Transformer model length limitations, enable the reporting of pathogenic capacity, protein importance, and genome pathogenic-related embeddings.

To train the model, we compiled the largest bacterial pathogenicity-annotated dataset to date, using genomic sequences from publicly available databases (Section 2, Methods). The pathogenic and non-pathogenic capacity subsets were built separately, so similar bacterial genomes with opposite phenotypes were not discarded. This was done to account for species that have both strains with and without pathogenic capacity. For the pathogenic genomes, any bacterial sequence isolated from an infection site was included, regardless of virulence or incidence of infection on humans. The non-pathogenic genomes had to meet all following three criteria: 1) not reported as pathogenic; 2) not isolated from a sick human; 3) defined as non-pathogenic. We defined four bacterial phenotypes that could fulfill the last requirement:

a) explicitly non-pathogenic bacteria; b) bacteria of the human microbiome; c) bacteria classified as probiotics or used in ingested products; d) extremophiles from environments not found in the human body. Thus, only strains categorically non-pathogenic or unable to live in the human body, or those repeatedly in contact with humans without causing disease, were selected. The final PathogenFinder2 dataset comprised 16,297 pathogenic and 4,882 nonpathogenic nonredundant genomes (a more detailed description of the process is included in Section 2, Methods).

## 2 Methods

### 2.1 PathogenFinder2 Dataset: Development and Test-NovelSpecies sets

The dataset for developing and testing PathogenFinder2 was constructed from public databases of annotated bacterial genomes. The dataset creation involved three major steps (Figure 2): 1) Collection of genomic entries annotated for pathogenic capacity; 2) Linking entries to genomic sequence repositories; 3) Homology reduction and splitting. Different approaches were used in the first step, depending on whether the bacteria had pathogenic capacity. For the pathogenic bacteria subset, data was collected from two databases: NCBIPathogen (https://www.ncbi.nlm.nih.gov/pathogens/, accessed October 2023) and ENAPathogen (https://www.pathogensportal.org, accessed October 2023). In NCBIPathogen, entries from the Isolates Browser were downloaded, and sequences with hosts labeled “human” or “homo sapiens” (excluding those with “not”) were selected. In ENAPathogen, bacterial assemblies from the Pathogen Sequences category were downloaded using ENA Advanced Search, and sequences with “human” or “homo sapiens” in the “host” or “host_scientific_name” columns, or taxonomy ID 9606 in the “host_tax_id” column, were selected. These databases were not available when other pathogenic capacity predictors were designed.

**Figure 2:**
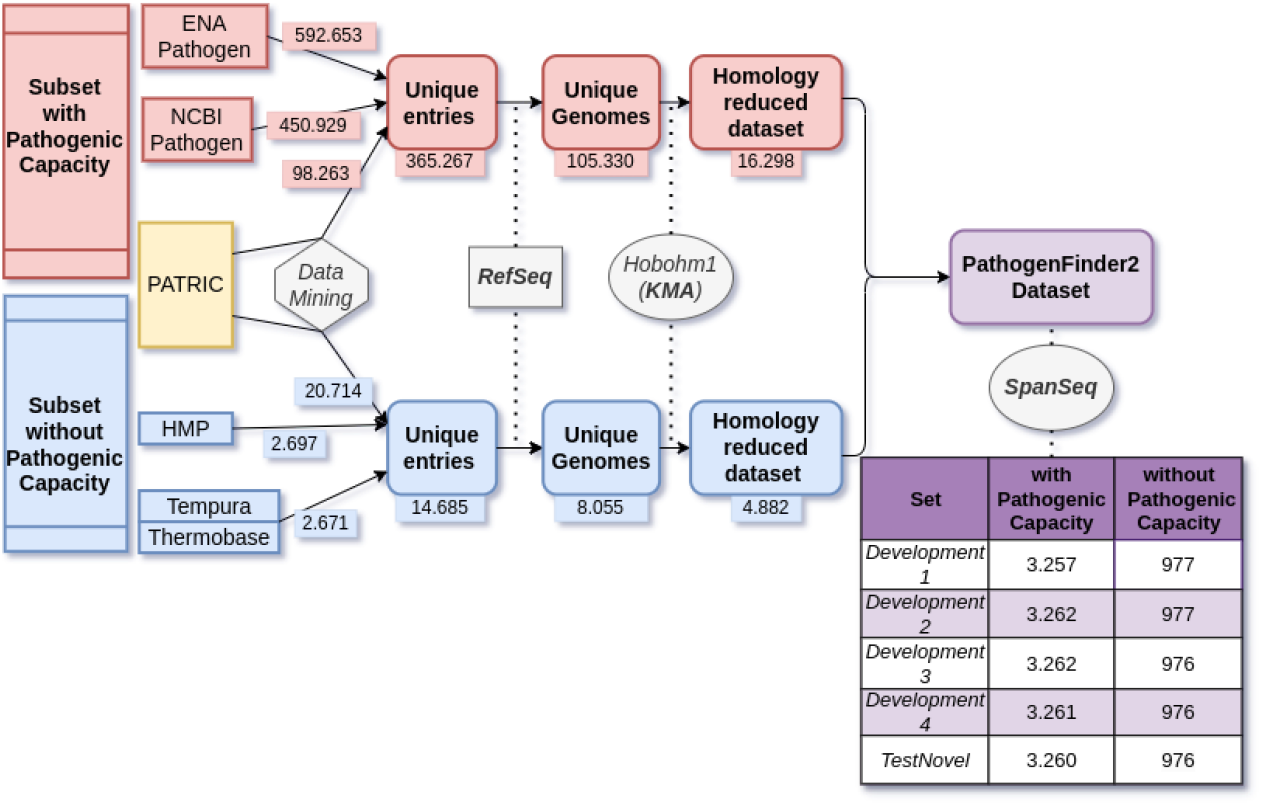
Overall scheme of the procedure to create PathogenFinder2 dataset. The scheme includes the amount of entries/genomes at each step, as well as the number of genomes with and without pathogenic capacity after homology splitting with SpanSeq. The development sets were used for hyperparameter optimization and training of the model.

To maximize data collection, the general annotated bacteria database PATRIC (accessed October 2023, https://www.bv-brc.org/) was also used. The database was filtered for keywords indicating pathogenic capacity, improving upon previous methods with data mining techniques. Six columns in PATRIC described disease-causing bacterial genomes: “Isolation,” “Source,” “Host Health,” “Other Clinical,” “Comments,” and “Additional Metadata.” The keyword extraction process involved:

- Tokenizing text in columns of interest
- Removing common English words (stopwords)
- Converting to lowercase and removing numeric/non-alphabetic tokens
- Creating a unique token collection
- Manually classifying tokens related to the desired phenotype (excluding species names)
- Create a unique stem collection from the tokens The manually selected vocabulary for pathogenic bacterial cases fell into four categories (Table 2): Pathogenic, Pathogenic-related, and Disease (1 & 2). The first category, “Pathogenic,” includes terms that unequivocally describe a bacterial isolate capable of causing harm to a host, such as “pathogenicity,” “virulence,” or “infection.” The second category, “Pathogenic-related,” encompasses terms describing generic events associated with pathogenic bacteria, such as “fatal,” “food-borne,” “water-borne,” “epidemic,” “damage,” and “outbreak.” The third and fourth categories consist of names of human diseases that are always or can be caused by bacteria, including some disease abbreviations. This vocabulary is attached in the Supplementary Material.

After creating the vocabulary, the same database was searched for bacteria with reported cases of human harm. Initially, only entries without contamination (as indicated in the “Host Health” and “Genome Quality Flags” columns), from the bacterial superkingdom, and with a human host (“Human,” “Humanus,” or “Homo sapiens” in the “Host Name,” “Host Common Name,” or “Isolation Source” columns) were selected. Subsequently, the extracted vocabulary was searched within the specified columns of the remaining entries. Each stem was searched, and selections were made only if the stem was not preceded by a negative term (e.g., “no,” “un,” or “vaccine”) and was not part of an organization name (e.g., “Center for Genomic Epidemiology,” which contains the stem “epidemi”). A schematic of the entire process is shown in Supplementary Material, Figure S8.

To create the subset of bacteria without pathogenic capacity, data was sourced from non-pathogenic specific databases and the general PATRIC database. For selecting bacterial extremophiles requiring conditions not present in human tissues, the entire Thermobase[19] database and a subset of the Tempura[49] database were used. Additionally, the first phase Human Microbiome Project[55] database was downloaded, as bacteria from healthy human microbiomes were considered mostly non-pathogenic. In PATRIC, a similar process for selecting pathogenic entries was followed to filter non-pathogenic bacteria. However, instead of keyword extraction, a manually curated dictionary was created with categories: Non-pathogenic, Microbiome, Probiotic, and Extremophile (Table 2).

The previous method for selecting entries of interest (without pathogenic capacity) was then applied, as shown in Supplementary Material, Figure S8. For “Probiotic” or “Extremophile” categories, bacteria did not need to have human hosts. An additional step was included to discard entries indicating unhealthy hosts or bacteria that spoil human-consumed products (the vocabulary defined as Unhealthy in Table 2).

Entries from various databases required a RefSeq[44] or GenBank[50] ID. Duplicates based on these IDs were removed, and if an ID had two different phenotypes, the non-pathogenic entries were discarded. The two subsets of genomes (with and without pathogenic capacity) were homology reduced separately using the Hobohm algorithm[26] in KMA[15], not allowing for sequences in the database with a higher query coverage (in k-mer space) than 90%.

To prevent data leakage when splitting the dataset into training and evaluation sets, we used SpanSeq[23], set to allow a maximum of 70% of k-mers shared between sequences to not be forced to go in the same set (Figure 2). This threshold was chosen in order to ensure genomes from the same species appeared in the same set, as shown in its article. The split was also stratified to ensure the minority label (without pathogenic capacity) was represented in all sets.

### 2.2 PathogenFinder2 Test-2024Strains

The dataset with new bacterial entries (Test-2024Strains) was created following the same procedure as the original dataset (Figure 2), without keyword extraction. Only entries submitted during 2024 were considered. Due to using the same vocabulary dictionaries from the original dataset, a manual review of the PATRIC entries was conducted, resulting in the deletion of 15 entries due to unclear pathogenic capacity. Since all entries were used for testing, SpanSeq was not required. We evaluated the performance of various pathogenic predictors on a collection of E. coli isolates from healthy and unhealthy humans[25]. This species was specifically chosen due to its common occurrence and potential for heterogeneous pathogenic capacity.

### 2.3 PathogenFinder2 Architecture

PathogenFinder2 employs a combination of methods to transform a genome into a suitable input for a neural network. Initially, Prodigal[28] predicts a collection of proteins from the genome, each of which is embedded using the ProtT5[20] protein language model. This language model was chosen for its performance in other tasks and the size of its embeddings. To prevent out-of-memory errors from long protein sequences, the maximum length of a protein was set to 9920 amino acids. Longer sequences are split into overlapping segments before embedding, ensuring PathogenFinder2 fits within a 24GB GPU for inference. The maximum length can be adjusted for different infrastructures, though this may affect model accuracy.

The embeddings are summed along the amino acid dimension, resulting in a vector of 1024 features for each protein. These vectors are stacked to form the 2D data input for a neural network ensemble, with dimensions P (number of predicted proteins) by 1024 features per position. The neural network architecture, inspired by vision models, is based on ConvNext[37] for image classification. It begins with an initial layer to reduce input features, followed by three ConvNext blocks. These blocks, with Layer Normalization[5], are suitable for PathogenFinder2’s input due to their ability to handle varying sequence lengths.

The ConvNext structure was modified to accept sequential inputs and maintain the sequence length (not the features) throughout the layers. The attention layer, based on the Bahdanau or additive attention mechanism[6], processes the sequence length without altering it. To provide structured information to the neural network, the number of predicted proteins is embedded using a positional encoding-like architecture and concatenated to the attention layer output. The way this information was introduced to the neural network (added, concatenated, or ignored) was selected through hyperparameter selection (Supplementary Material, Table S2). This concatenated output is fed into a sigmoid layer for classification and serves as the bacterial genome embedding.

PathogenFinder2’s neural network model is an ensemble of four networks, each trained on different training/validation splits. The final prediction is the mean output of the four networks.

### 2.4 PathogenFinder2 hyperparameter optimization and training

When designing the neural network architecture, parameters such as weight initialization, kernel size, batch size, and optimizer were initially chosen based on preliminary studies and literature. However, seven parameters required hyperparameter optimization (Supplementary Mateiral, Table S2). We employed the Tree-structured Parzen Estimator (TPE)[9] algorithm from Optuna[2], aiming to maximize the Matthews Correlation Coefficient (MCC)[38] of the neural network ensemble. Each neural network was trained for 60 epochs without pruning, selecting the best MCC value of each neural network using a 4-fold cross-validation approach. Only the last 50 epochs were considered, to avoid selecting initial epochs where training is more unstable. Despite aiming for a generalizable neural structure, the hyperparameter search yielded minimal improvement over initial guesses. We conducted 50 guesses controlled by the TPE algorithm, followed by 10 additional guesses based on observed results.

During both hyperparameter optimization and final training, we used bucketing techniques[32] and maintained a fixed class proportion per batch to address class imbalance. The RAdam optimizer[36] and a learning rate scheduler, which decreases the learning rate if the loss increases during training, were utilized, with an initial warm-up period. For final training, each neural network was trained for 200 epochs, selecting the weights from the best-performing epoch on the validation set. The neural network models were implemented in PyTorch[47], and the models were trained (including hyperparameter optimization) and tested on GPUs Tesla NVidia A100 with 40 GB HBMI2, with CPUs AMD Epyc 7H12.

### 2.5 Other Models: Previous Pathogen predictors

We tested the previous protein-based bacterial pathogen predictors PathogenFinder1, BacPacs and WSPC, as well as the read-based predictor DeePac on the Test-NovelSpecies and Test-2024Strains. Prodigal was run with standard setup to produce the protein predictions for PathogenFinder1 and BacPacs methods. For PathogenFinder1, the same code that runs behind the WebServer (https://cge.food.dtu.dk/services/PathogenFinder/) was used (provided by the department that hosts it), with the standard setup and with the “All” database (database without any phylum specified). The probability value given by the algorithm was used to build the ROC and calibration curves. For BacPacs, we followed the instructions of the section “Predicting data using bacpacs pre-trained model” available from their repository (https://github.com/barashe/bacpacs). As BacPacs only provides a label as output, we used the probability estimation for OvR logistic regression (_predict_proba_lr) of the linearSVC that the model uses for predicting pathogenicity. The probabilities are computed as: 1*/*(1 + *exp*(*− decisionfunction*(*X*)). For WSPC, we followed the instructions available on their repository (https://github.com/shakedna1/wspc_rep) using their existing model. We obtained the PATRIC Global Protein Families (PGFams), required by the model, through PATRIC’s Genome Annotation Service (https://www.bv-brc.org/app/Annotation) as described in the repository. As we have not given any information to the other models about the taxonomy of the test set, we set the Taxonomy ID to 2 (bacteria). For DeePac, we followed the instructions described in their repository (https://gitlab.com/rki_bioinformatics/DeePaC). As it requires the input to be reads, we used their method to produce pseudoreads (deepac gwpa fragment), before predicting bacterial pathogenicity. BacPacs, WSPC and PathogenFinder1 models were tested on a Linux system on Intel(R) Xeon(R) CPU E7-8870 v4 @ 2.10GHz. DeePac was tested on NVIDIA Tesla V100 SXM2 with 32 GB GPUs with PCIE.

### 2.6 Other Models: Baseline Pathogen predictors

The baseline method Kmer-Patho is an adaptation of the KmerFinder taxonomy detection method, which examines the number of co-occurring k-mers[24]. We used the code available in the repository (https://bitbucket.org/genomicepidemiology/kmerfinder.git). To shape the PathogenFinder2 dataset so it can be used by our tailored KmerFinder, we indexed the development subset (without overlap with Test-NovelSpecies) using KMA with “*kmaindex ™ i < in*.*fna > ™ o < OUT > ™ SparseATG*”, as used in the database behind the webserver https://cge.food.dtu.dk/services/KmerFinder/. Unlike the species predictor, each sequence was only defined by its pathogenic capacity. The prediction was then done taking the phenotype of the closest sequence. To provide a continuous output (in order to produce the ROC and Calibration curves), the query coverage of the hit on the database was used. The other baseline method created was a version of PathogenFinder1 with the new database. We followed the procedure described by the authors and that is implemented on the code behind the webserver https://cge.food.dtu.dk/services/PathogenFinder. Due to the significant increase in the protein database size, CD-Hit[34] was replaced by the state-of-the-art clustering method MMSeqs2[52] for building the database (clustering proteins, using the “linclust” clustering strategy), as well as for searching for matches (using the “search” module). Both baseline models were created and tested on a Linux system with an Intel Xeon E7-8870 v4 @ 2.10GHz CPU

## 3 Results

### 3.1 PathogenFinder2 excels in predicting novel pathogens

Our novel model, PathogenFinder2, achieved a higher Matthews correlation coefficient (MCC) compared to both the baseline models trained with the new dataset (PathogenFinder1DB2, KmerPatho) and previous state-of-the-art models (PathogenFinder1[16], WSPC[42], BacPacs[7], DeePac[8]) on new species (Test-NovelSpecies). For the dataset allowing for leakage (similarity between training and testing allowed, Test-2024Strains), the MCC for PathogenFinder2 remained higher than for PathogenFinder1DB2; however, this difference was not statistically significant according to the 95% confidence interval (CI, i.e., within ±1.96 standard errors; Figure 3(a)). PathogenFinder2 also demonstrated a superior ROC curve and higher AUC compared to other methods in Test-NovelSpecies (Figure 3(b)). On the test set permitting similarity, PathogenFinder2 exhibited metrics comparable to PathogenFinder1DB2, though it had a higher true positive rate when the false positive rate was near zero. The species-based baseline method, KmerPatho, performed significantly worse than both PathogenFinder2 and PathogenFinder1DB2 in terms of MCC and AUC, especially when there was no species overlap between its database and the test set, in which case it was also outperformed by previous pathogenic predictor methods.

**Figure 3:**
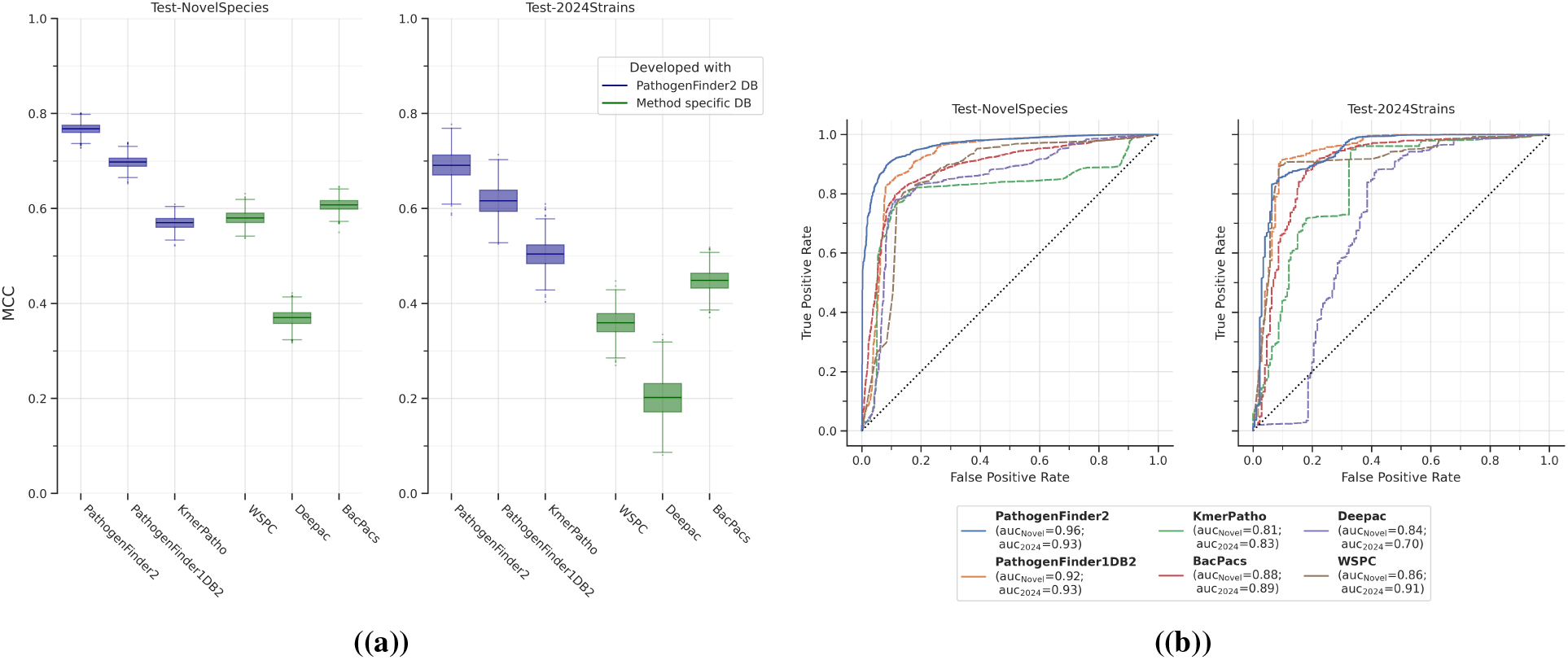
Performance measures of PathogenFinder2, Baseline Methods, and Previous State-of-the-Art Methods. (a) Box plots showing the 5 quartiles of the MCC values from bootstrapping on the different test sets. (b) ROC curves with AUC in the legend. On the left panels, Test-NovelSpecies (3,310 pathogenic, 976 non-pathogenic), with bacterial species not included in training PathogenFinder2 or baseline models. On the right panels, Test-2024Strains (3,704 pathogenic, 176 non-pathogenic) with bacteria annotated a year after PathogenFinder2 dataset creation. Besides PathogenFinder2, we evaluated two baseline models trained with the PathogenFinder2 database, PathogenFinder1DB2 (PathogenFinder1 trained with the new dataset) and KmerPatho (a method based on the k-mer based species predictor KmerFinder[24]), and four previous state-of-the-art pathogen predictors. The methods were executed using their pretrained configuration or database, as specified in their repositories. Further details are provided in the Section 2, Methods. The original PathogenFinder1 failed to generate any result for 1,660 genomes in Test-NovelSpecies and 2,189 in Test-2024Strains, so it was excluded from the comparison.

In terms of sensitivity, PathogenFinder2 was only surpassed by DeePac on Test-NovelSpecies (Supplementary Material 1, Figure S1a). However, DeePac exhibited very low specificity (Supplementary Material 1, Figure S1b) due to a high rate of false positives (Supplementary Material 1, Figure S3). BacPacs was the only method with higher specificity than PathogenFinder2, although this difference was again not statistically significant. Additionally, PathogenFinder2 showed a remarkably well-calibrated output for a deep learning model, particularly in the more extreme output regions (0-0.3 and 0.8-1 in Supplementary Material 1, Figure S2).

### 3.2 Attention highlights proteins related to pathogenicity

We utilized the weights provided by the attention module to report protein importance in the pathogenic capacity prediction. The top highlighted proteins by the attention module can be mapped to a protein database, such as UniRef50[53], as we did for our Test-NovelSpecies and Test-2024Strains sets (Supplementary Material 2). In the bacteria predicted with pathogenic capacity, most proteins fell into various virulence factor categories described in previous literature[35] (Supplementary Material 1, Table S1), such as proteins involved in adhesion to host tissues, intracellular invasion and toxins, as well as metabolic pathways such as glucose and iron metabolism, and anaerobic lifestyle. Besides, there was a significant number of genes related to genetic mobility, which can point to the involvement of mobile pathogenicity elements. In the bacteria predicted without pathogenic capacity from our Test-NovelSpecies and Test-2024Strains, most proteins lacked an annotated function. But among the proteins annotated with a function, we found antimicrobial producing genes, protein antidotes, glucosidases for degrading lactose and cellulose, and proteins involved in the shikimate pathway and chlorophyll production.

### 3.3 The Bacterial Pathogen Capacity landscape

We also offer an initial overview of the human bacterial pathogen landscape with the bacteria with pathogenic capacity of our dataset (Figure 4). We use PathogenFinder2’s capacity of embedding a genome into a fixed-length vector based on its pathogenic capacity. As expected, there appears to be correlation between closeness and the phylogenetic proximity of the species. However, we also observe patterns indicating that the embeddings contain additional information, such as the site of infection. For example, *Campylobacter concisus* is distinctly separated from other *Campylobacter* species that infect the gastrointestinal tract, but closer to skin flora. Similarly, *Streptococcus pneumoniae* is grouped with bacteria that cause lung infections rather than with other *Streptococcus* species (including *S. pyogenes* and *S. agalactiae*), as part of the oral flora.

**Figure 4:**
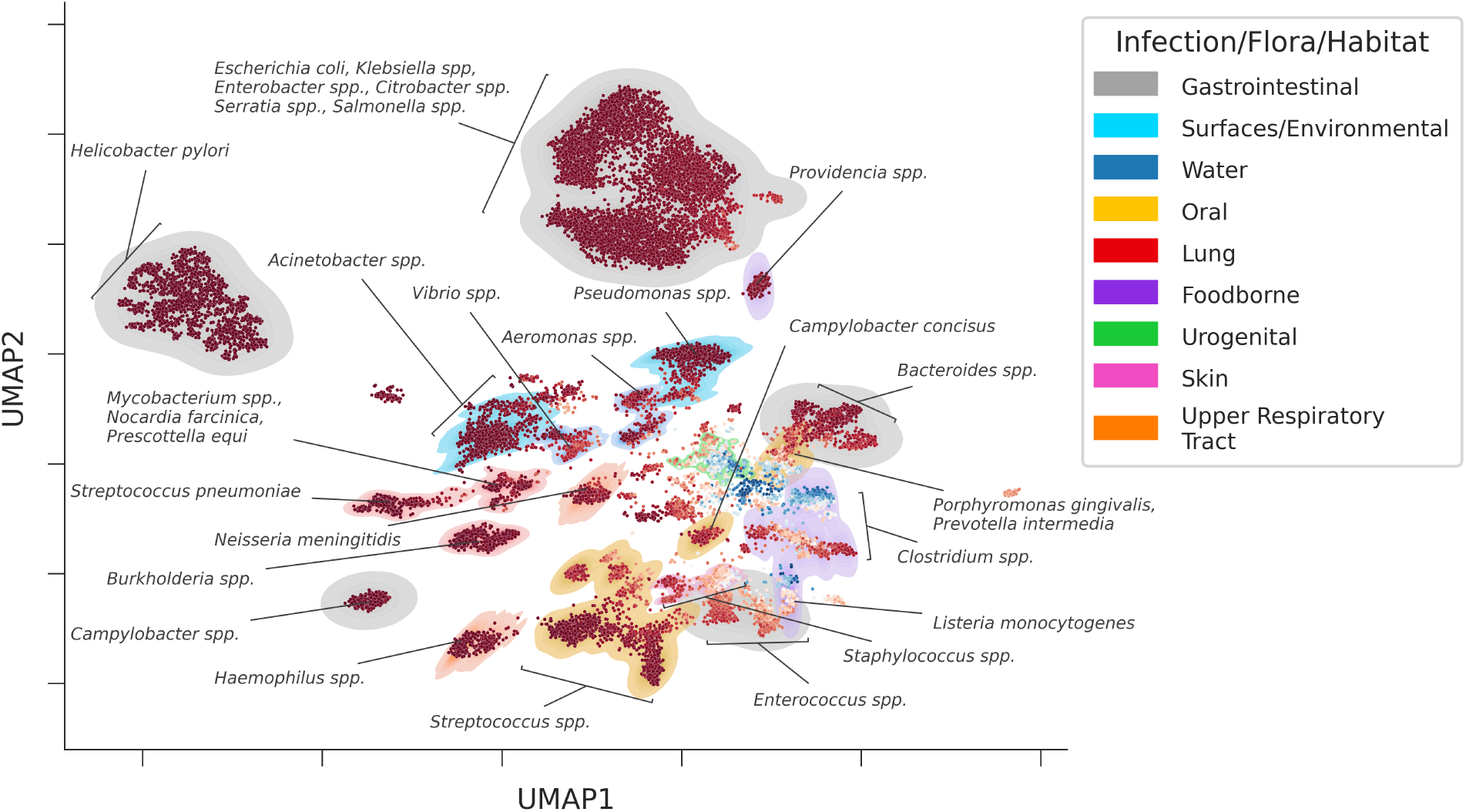
Projection of the embeddings of all bacterial sequences in the PathogenFinder2 database with pathogenic capacity, illustrating the Pathogen Capacity Landscape. Each dot represents a bacterial sequence that was embedded using PathogenFinder2 and plotted in a UMAP[40] of two dimensions. The dots are colored depending on the PathogenFinder2 pathogenic capacity prediction, with red being positive and blue negative The distribution representing the infection, flora or habitat of the bacteria was drawn with a kernel density estimation, and colored through a process of clustering with HDBScan[39] (as shown in the Figure S4 in Supplementary Material) and manual annotation based on bibliography. Some species and genera of interest are indicated with arrows.

Common oral pathogens such as *Porphyromonas gingivalis, Prevotella intermedia*, and *C. concisus* appear closely related to foodborne species from the *Clostridium* genus, like *C. perfringens, C. difficile*, and *C. botulinum*. These oral pathogens surround a miscellaneous group of bacteria known for degrading and fermenting biological material (such as the *Bacteroides* genus, part of the gut microbiota but isolated from infections in soft tissues and bacteremia), suggesting metabolic similarities between oral and foodborne pathogens rather than with oral flora. The latter, mainly part of the *Streptococcus* genus, have been reported to cause infections in other body parts, such as bacteremia or endocarditis. They are located next to species colonizing the upper respiratory tract from the *Haemophilus* genus (like *H. influenzae* or *H. haemolyticus*), which also cluster together with *Pasteurella multocida*. Although *P. multocida* is not part of the human upper respiratory tract flora, it is of certain animals and can be transmitted through their bites[4]. Another upper respiratory tract flora member, *Neisseria meningitidis*, appears close to lung infection pathogens. Although rare, *N. meningitidis* can cause pneumonia[22], which may explain this distribution.

The embedded representation of the bacterial proteome does not always correlate with the site of infection, but rather their habitat. For instance, *Vibrio* species like *V. parahaemolyticus* and *V. cholerae* appear closely to common nosocomial pathogens such as *Acinetobacter spp*. Or the known foodborne pathogen *Listeria monocytogenes* is placed alongside *Enterococcus casseliflavus*, a species that has only been seen to cause diseases in immunocompromised patients[58].

However, the capacity of some strains of this species to prevent *L. monocytogenes* infection by competing with their colonization using bacteriocins[48], could explain their proximity in the landscape. Moreover, *Klebsiella pneumoniae*, part of the gut microbiota but able to infect pulmonary tissues, appears in a cluster alongside gastrointestinal pathogens. The dimensions and spread of this cluster could suggest genomic similarity, due to, for example, the exchange of mobile elements[31]. Other clusters of bacteria that cause gastrointestinal diseases, like *Helicobacter pylori* or *Campylobacter spp*., show very different conformations. While the cluster of the latter, with two species (*C. jejuni* and *C. coli*), forms a dense cluster, the one with only one species, *H. pylori*, appears dispersed, likely due to the notable genomic variability of the species[1].

### 3.4 Practical case: bacterial pathogenicity prediction on metagenomic samples

PathogenFinder2 was employed to analyze 2,739 metagenomic assembled genomes (MAGs), many of which were completely novel taxa, from a previous study on sewage bacterial communities[30]. The analysis predicted that 1,839 bacteria lacked pathogenic capacity, while 370 were potentially capable of infecting humans under certain conditions. The remaining 530 MAGs were classified as Uncertain due to the neural network predictions being close to the 0.5 threshold. Using the embeddings of the genomes, we identified seven groups of potential pathogens (Supplementary Material, Figure S5 and S6). Each cluster was defined at the genus or family level (Table 1), including known pathogenic groups such as *Pseudomonas, Enterobacteriaceae*, and *Streptococcus*, as well as *Brachymonas*, which have not yet been reported to cause infections.

**Table 1:** Keywords for data mining of PATRIC. Categories “Pathogenic, Pathogenic-related”, “Disease” and “Unhealthy” were created using the method of keyword extraction. Categories “Non pathogenic”, “Probiotic”, and “Extremophile” were created manually. The section “Unhealthy” is not used as a basis for creating any subset of the database, but to control that the entries of the non-pathogenic subset were coming from healthy humans (if not from probiotics or extremophiles).

| Category | Examples (Stems) | Used In |
| --- | --- | --- |
| Pathogenic | pathogen-, virulen-, infec-, superbug-, ... | Pathogenic Subset |
| Pathogenic related | fatal-, borne-, epidemi-, damag-, toxi-, outbreak-, endemi-, ... | Pathogenic Subset |
| Disease (1) | necrotis-, bacteremi-, pneumoni-, botulism-, diphtery-, ... | Pathogenic Subset |
| Disease (2) | gangrene-, diarrheal-, polyp-, encephalitis-, larygnitis-, ... | Pathogenic Subset |
| Unhealthy | accident, amputation, obstruction, subacute, bleed-, | - |
| Non Pathogenic | non-patho-, volunteer-, commensal-, reference-, donor-, ... | Non-Pathogenic Subset |
| Microbiome | flora, microbiom-, gut-, inhabit-, microbiot-, ... | Non-Pathogenic Subset |
| Probiotic | probiotic, novel food, cheese, kefir, fermented, wine, ... | Non-Pathogenic Subset |
| Extremophile | hyperpiezophile, piezophile, termophile, cryophile, ... | Non-Pathogenic Subset |

**Table 2.**
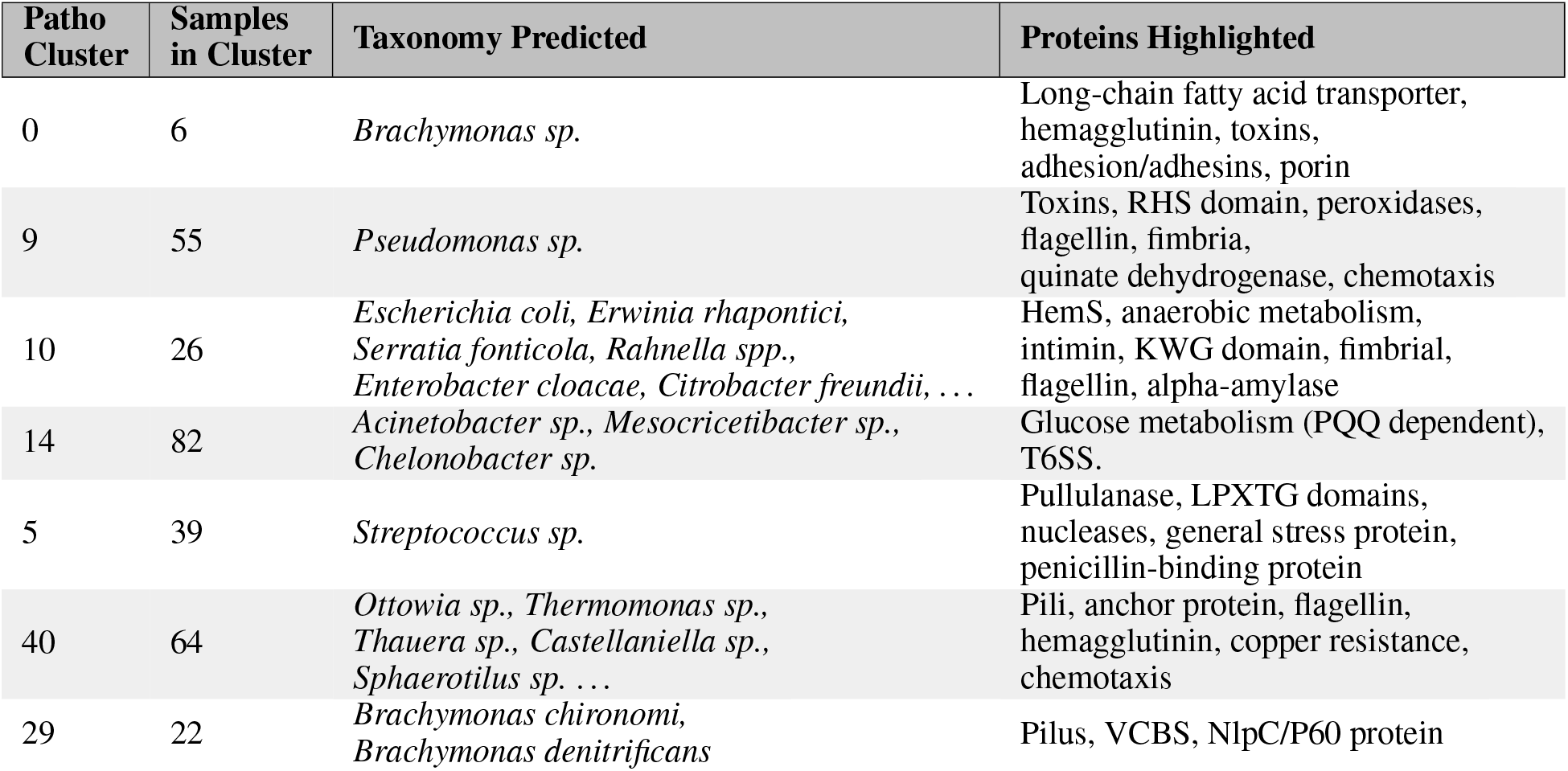
Pathogenic Clusters in Metagenomic Samples. Pathogenic clusters identified in metagenomic samples (Supplementary Material, Figure S3) show their major predicted taxonomy and proteins highlighted by the attention vector. Mapping these clusters onto the phylogenetic tree from study [30] revealed that pathogenic phenotypes were not consistently homogeneous within each clade (Supplementary Material, Figure S4). Examination of the protein content highlighted by the attention layer for each cluster reveals known virulence factors. Each cluster exhibits distinct protein profiles, helping in their definition; for example, Cluster 5 may contain intracellular pathogens.

## 4 Discussion

PathogenFinder2 does not aim to predict bacterial pathogenicity, as the interaction outcome between bacteria and humans also depends on the latter[41, 13]. Instead, we redefined the objective to predict bacterial pathogenic capacity, which is solely determined by the bacteria. The dataset used to train the model was designed to reflect this definition, as it would shape the deep learning model accordingly. Genomes were classified as “pathogenic” if they had caused any human infections, regardless of virulence, host conditions, or incidence. This broader definition should be considered when using PathogenFinder2, as it may predict bacteria with pathogenic capacity that might require specific host conditions to cause harm. For instance, there is no difference in the predictions of PathogenFinder2 (and other pathogen predictors used in this study, Supplementary Material Figure S7) on *E. coli* strains found in diarrheal and non-diarrheal patients[25]. Naturally, the pathogenic capacity predictors might not be fine-grained enough to detect minimal genetic differences between pathogenic and nonpathogenic *E. coli*[33]. But, being all predicted with pathogenic capacity, the difference on the interaction outcome might not be due to different pathogenic capacity, but for reasons only reliant on the host.

Additionally, labeling bacteria as pathogenic based on their isolation from an infection site carries the risk of mislabeling non-pathogenic microbiota, as they might be present in the affected area but not causing harm. In fact, the presence of noise in bacterial database annotations is unavoidable. Previous attempts to create pathogen predictors have tried to minimize this issue by arbitrarily selecting one phenotype for some heterogeneous species[42, 17, 8]. However, this approach does not acknowledge the existence of species with heterogeneous pathogenic capacity among their strains. To avoid simplifying the neural network’s objective to a mere species predictor, we relied on genomic annotations provided by researchers uploading genomes to publicly available databases, allowing similar genomes with different phenotypes. Although this might introduce some noise into the PathogenFinder2 dataset, we believe the model has overcome these noisy labels. For example, the sequences predominantly predicted as non-pathogenic shown in the Pathogen Landscape (Figure 4) by PathogenFinder2 mainly consist of bacteria, such as *Bacteroides* or *Lactobacillus*, found in the human microbiome that populates common infection sites of other bacteria.

Another consideration when working with bacterial pathogenicity is the potential bias in our dataset’s subset of bacteria without pathogenic capacity. Besides extremophiles unable to live in the human body, we required that these bacteria were frequently in contact with humans without reported infections. This might have skewed the subset and may not represent reality. This compromise is unavoidable, as creating new data would require testing bacterial pathogenic capacity in humans. Additionally, the ratio of bacteria with and without pathogenic capacity in the database may not reflect real ecosystems. For instance, only 13% of bacterial genomes isolated from sewage[30] were predicted to have pathogenic capacity, compared to 76.9% in our training dataset. Although we cannot generalize the ratio found in sewage to other ecosystems, it demonstrates our model’s ability to learn beyond the dataset imbalance.

Our model outperforms the previous state-of-the-art pathogenicity predictors on novel species, with balance between specificity and sensitivity in its predictions. This is particularly noteworthy given that PathogenFinder2 was trained without exposure to these species, unlike the previous models that may have overlapping species between their training sets and our test sets. PathogenFinder2 effectively overcomes data-related challenges that our baseline models struggle with, showing that our model’s success can not only be attributed to the creation of the largest bacterial pathogen dataset created, but also to the model’s architecture. For instance, PathogenFinder1DB2 (PathogenFinder1 trained with the new dataset), like any alignment-based model, suffers from lower accuracy on unseen species due to dataset incompleteness, in particular when predicting the under-represented label (non-pathogenic). Additionally, KmerPatho underperforms compared not only to PathogenFinder2, but also against older models, particularly on unseen species, indicating that pathogenic capacity prediction cannot be achieved using species prediction models.

PathogenFinder2 combines the advantages of not requiring a tailored database of proteins related to pathogenic capacity (as in previous read-based models), while still identifying proteins of interest for its predictions (as in previous protein-based models). Although attention scores as explanations for pathogenic capacity pose challenges[56, 29], they offer valuable insights for surveillance programs and researchers (Table 1). Moreover, the lack of causality in attention weights is not necessarily a drawback, as it might provide an insight to new virulence factors and metabolic infectious pathways not found by more traditional methods.

Iron acquisition proteins[14] and several proteins related to glucose intake and metabolism[46], particularly PQQ-dependent and anaerobic condition-related proteins, are constantly highlighted by the model. Enzymes involved in genetic stability appear repeatedly and exclusively in pathogenic bacteria, suggesting the importance of genomic variability and mutation rates for bacterial adaptation to hosts, as previously found for Salmonella enterica and E. coli[45, 54]. Identifying the specific functions of these proteins and metabolic pathways can provide crucial insights into infectious diseases and pave the way for the development of broad-spectrum antimicrobials.

PathogenFinder2 can also be used to create an embedding of a bacterium related to its pathogenic capacity. This has allowed us to produce the initial mapping of the Pathogenic Capacity Landscape, providing insights into the diverse mechanisms by which bacteria inflict harm on human hosts. Additionally, it can aid in the study of rare and under-researched pathogens, as well as interactions between bacterial species. PathogenFinder2’s embeddings encapsulate information regarding the infection site and habitat of pathogenic bacteria, among other features, although that information was not part of the training data. Future directions for our model may include incorporating this information when predicting pathogenic capacity, offering a more realistic and informative output regarding the threat certain bacteria pose to humans.

The aim of PathogenFinder2 is to provide a tool for surveillance of novel pathogens and the study of human infections. With the resourceful output of our model, available for local use or through our portal (http://genepi.food.dtu.dk/pathogenfinder2), we believe it will enable more targeted early detection of future pathogens, as well as the development of treatment strategies and vaccinations even before infections are observed, as demonstrated on the analysis of the MAGs from sewage[30]. Moreover, to the best of our knowledge, represents the first successful attempt to utilize pLMs for the prediction of whole genomes as input, and the analyses can now be expanded also to viruses and parasites, as well as animal and plant pathogens.

## 5 Acknowledgements

European Union’s Horizon 2020 research and innovation program under VEO grant [874735].

The authors gratefully acknowledge the HPC RIVR consortium (https://www.hpc-rivr.si) and EuroHPC JU (https://eurohpc-ju.europa.eu/) for funding this research by providing computing resources of the HPC system Vega at the Institute of Information Science (https://www.izum.si/en/home/).

