## Supplementary Material 1 for "Whole-genome prediction of bacterial pathogenic capacity on novel bacteria using protein language models, with PathogenFinder2"

*1. Research Group for Genomic Epidemiology, National Food Institute, Technical University of Denmark, Kongens Lyngby, Denmark; 2. Informatics and Predictive Sciences Research, Bristol Myers Squibb Company, Sevilla, Spain; 3. Bioinformatics, Health Tech, Technical University of Denmark, Kongens Lyngby, Denmark; 4. Rostlab, Department of Bioinformatics and Computational Biology, Technical University of Munich, Munich, Germany*

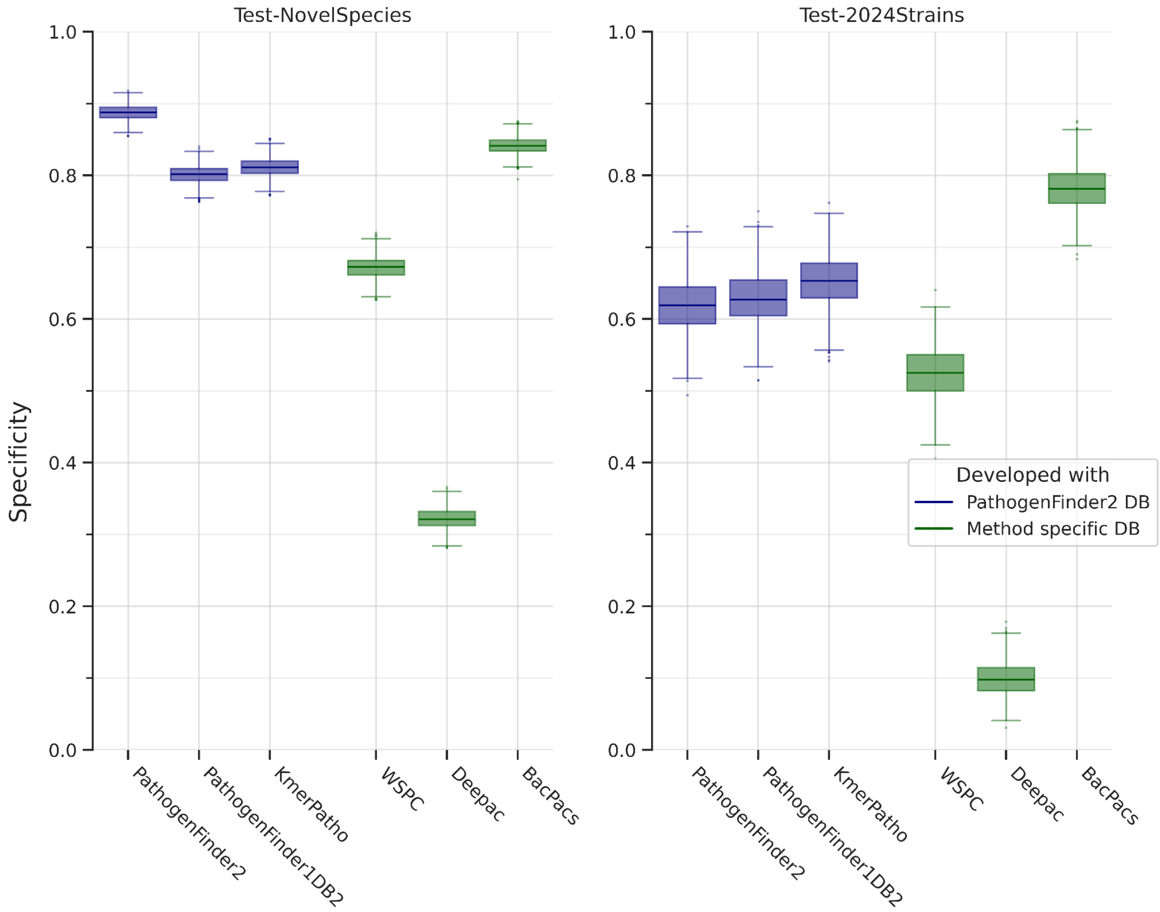
Test-NovelSpecies
Test-2024Strains
Specificity
1.0
0.8
0.6
0.4
0.2
0.0
PathogenFinder2
PathogenFinder1DB2
KmerPatho
WSPC
Deepac
BacPacs
Developed with
PathogenFinder2 DB
Method specific DB

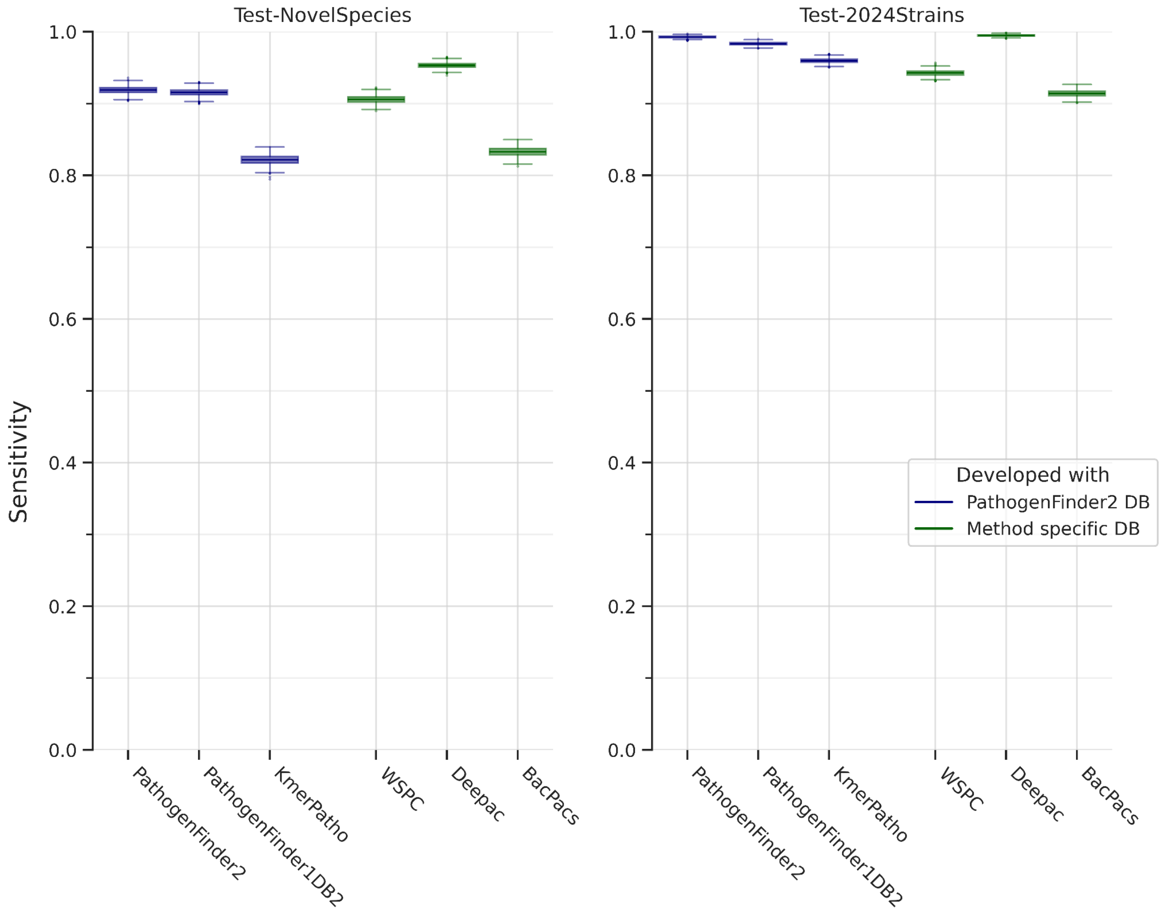


**Figure S1: Specificity and Sensitivity on the Test-NovelSpecies and Test-2024Strains from the PathogenFinder2, previous state-of-the-art methods and baselines.**

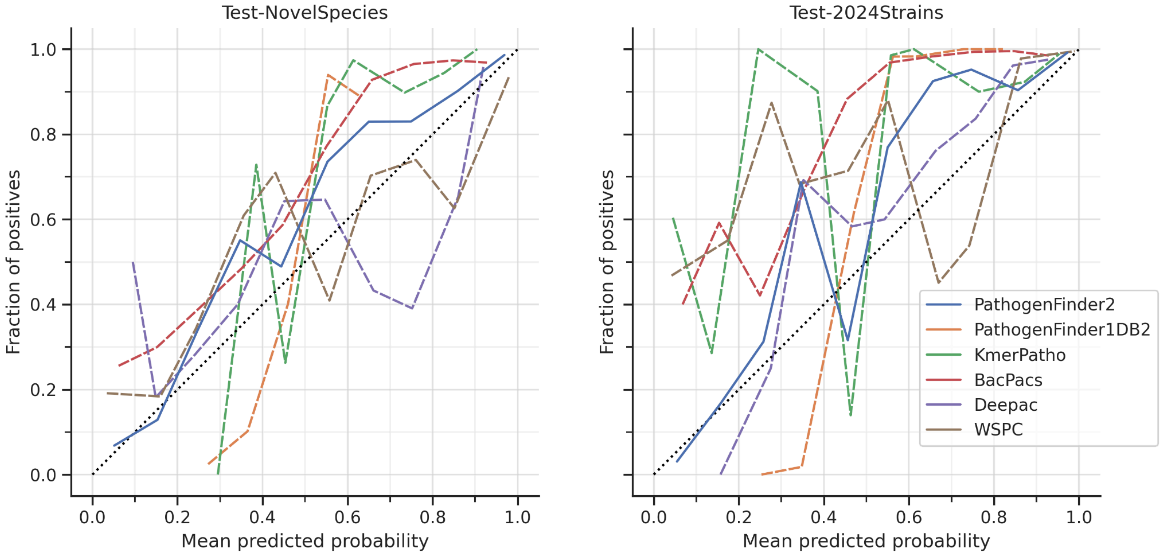


**Figure S2. Calibration curves on the Test-NovelSpecies and Test-2024Strains from**

**the PathogenFinder2, previous state-of-the-art methods and baselines.**

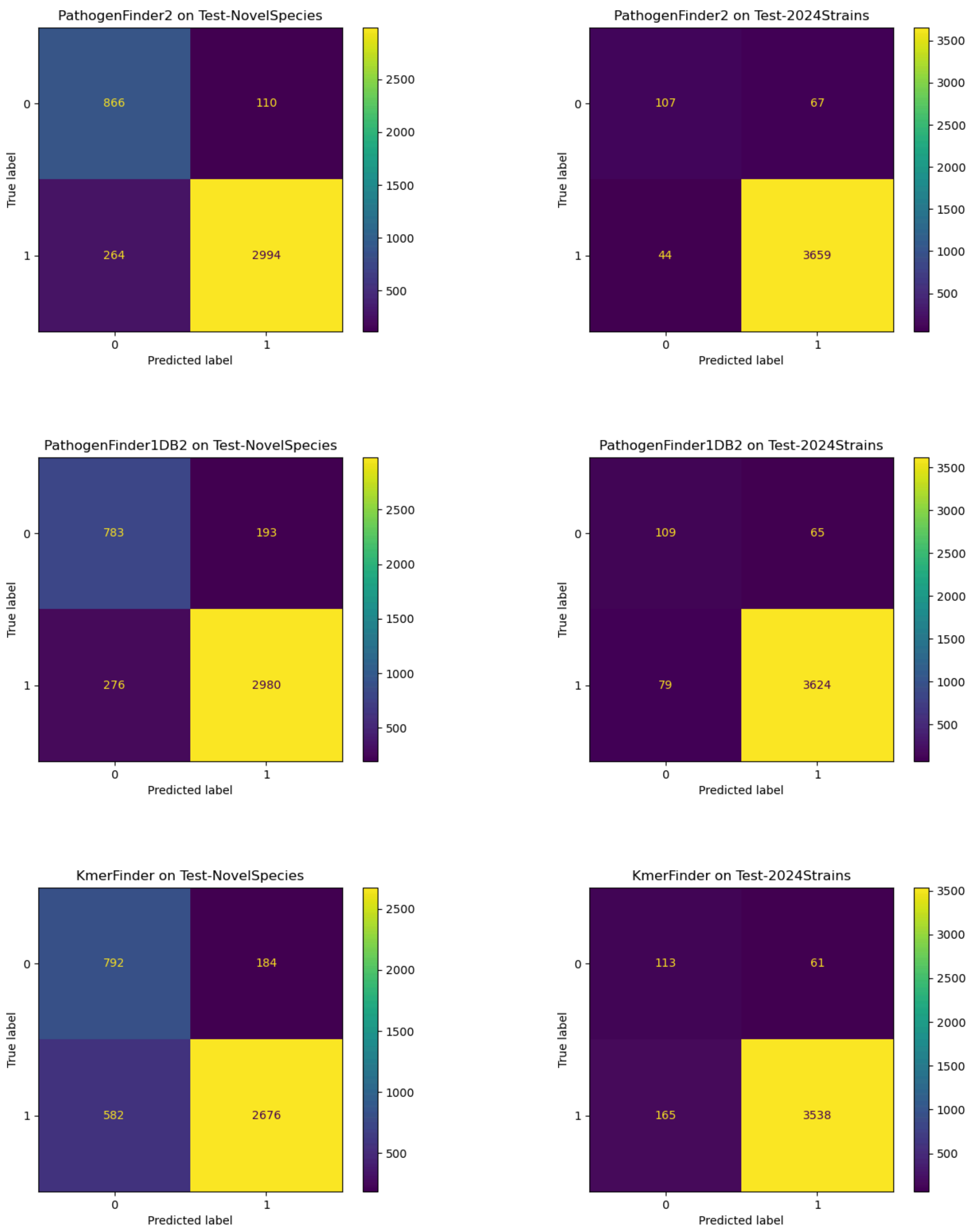

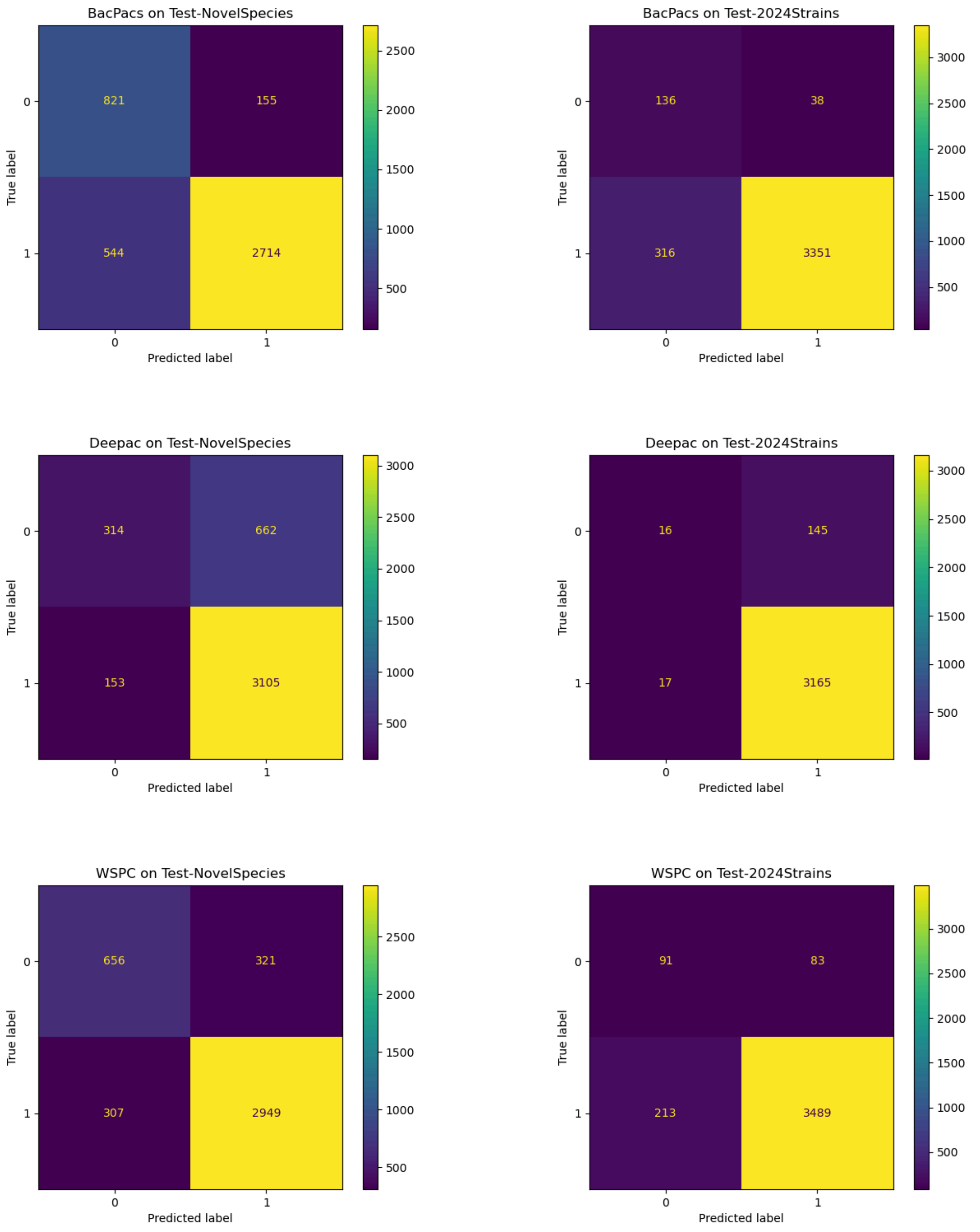


**Figure S3. Confusion tables on the Test-NovelSpecies and Test-2024Strains from the PathogenFinder2, previous state-of-the-art methods and baselines.**

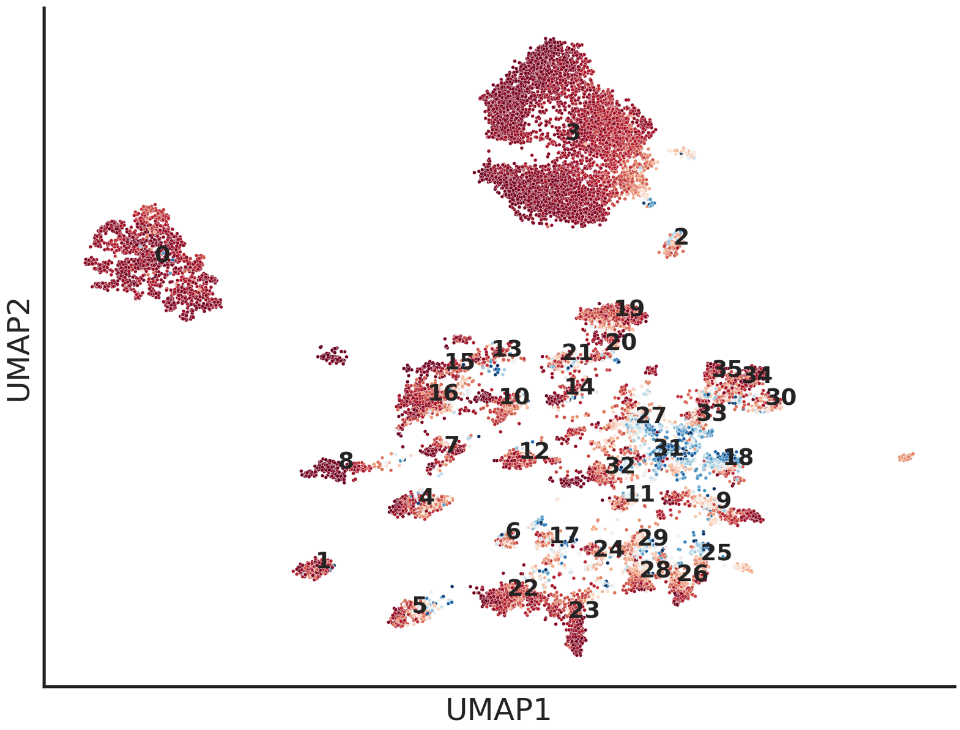


**Figure S4: Predictions of PathogenFinder2 on the pathogenic subset of the PathogenFinder2 dataset.**
The colors indicate the output of the sigmoid of the neural network model: from red (1, Pathogenic) to blue (0, Non-pathogenic). The UMAP is done on the embeddings produced by PathogenFinder2. The numbers indicate the clusters created with HDBScan, which can be mapped to the list of clusters in the Table S3, to map species on the Bacterial Pathogen Landscape.

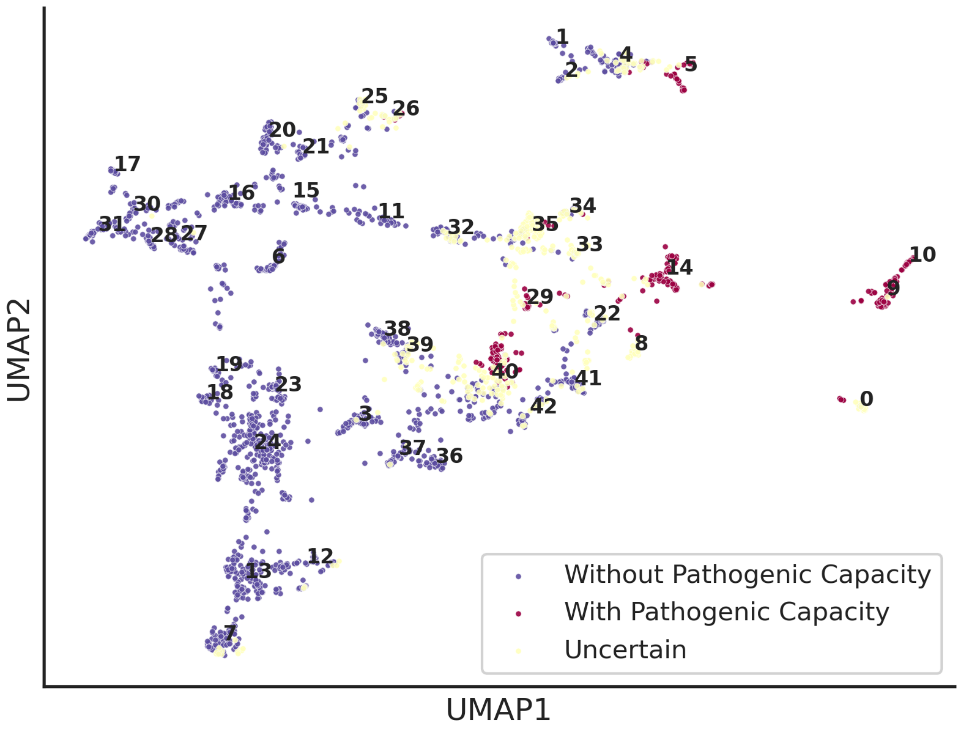


**Figure S5: Distribution of PathogenFinder2 embeddings of the MAG sequences from (Jespersen et al., 2023)**
In color is the prediction done by the model. The numbers indicate the cluster that the sequences below belong to, done with HDBScan.

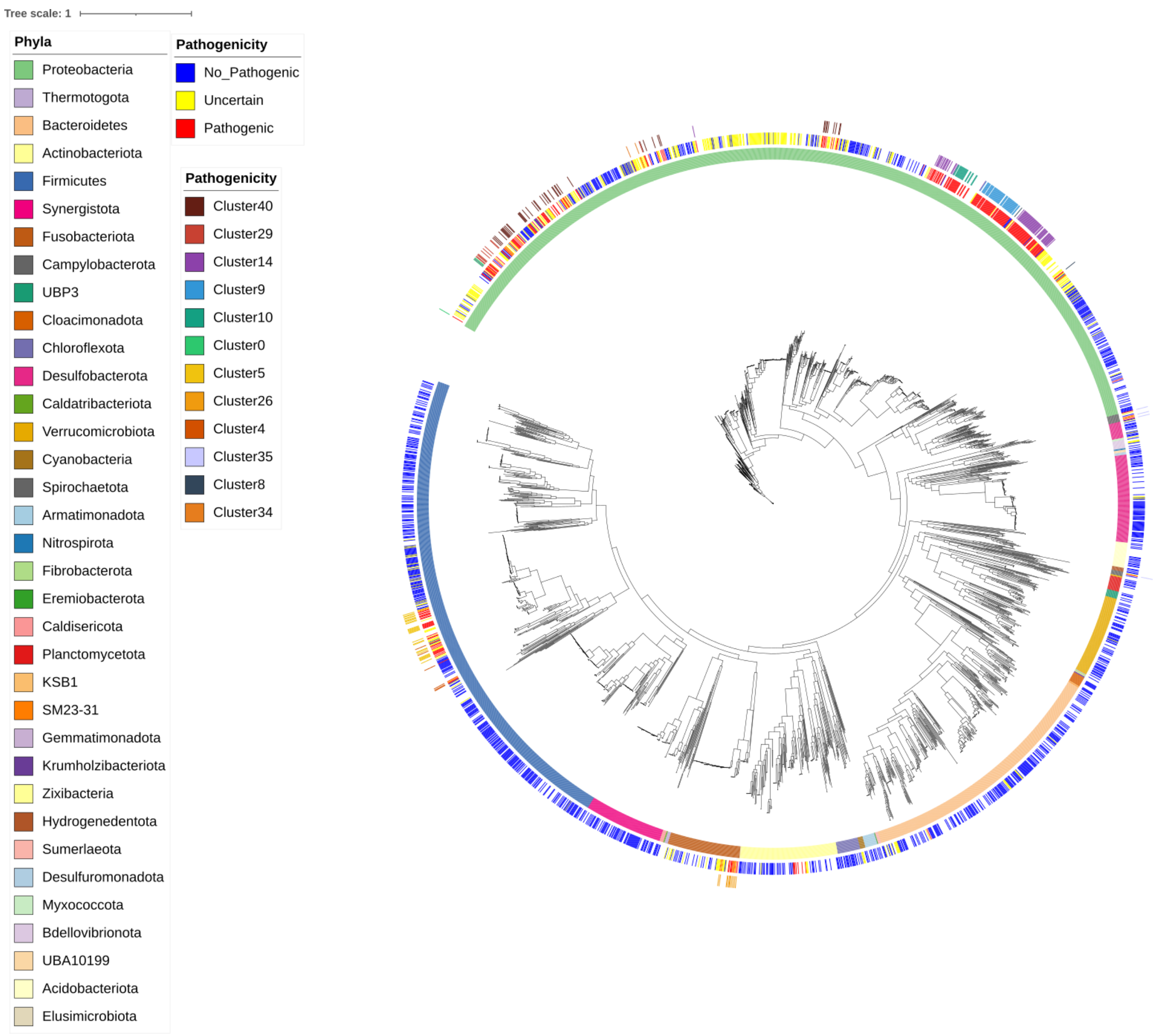


**Figure S6: Taxonomy of the MAGs from (Jespersen et al., 2023).**
The first circle is the phyla predicted by Jespersen et al., the second the predictions of PathogenFinder2, while the third circle highlights clusters with pathogens indicated in Figure S5.

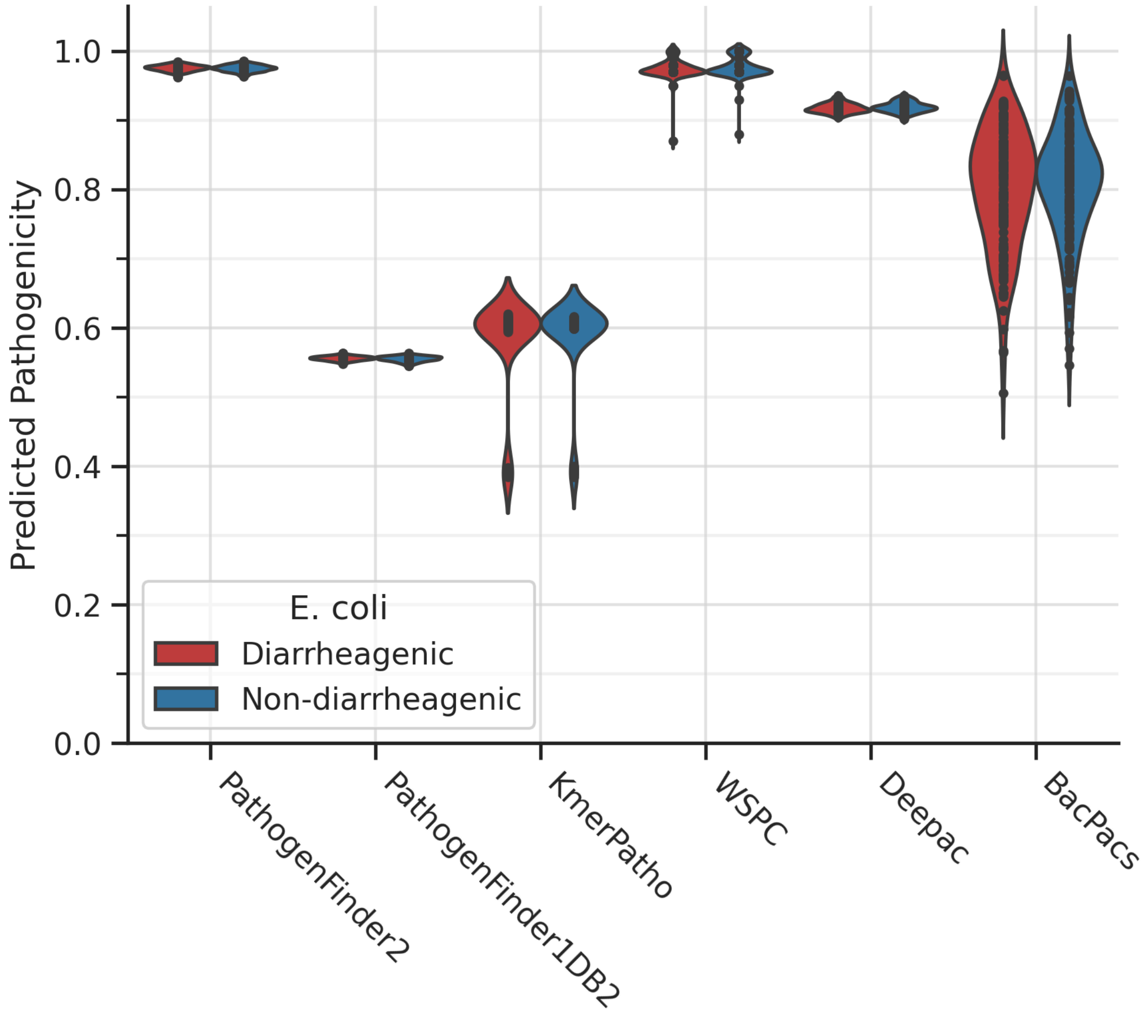


**Figure S7: Predictions of the different pathogenic capacity predictors on the E. coli samples collected from diarrheagenic and non-diarrheagenic samples from (Hazen et al., 2023).**

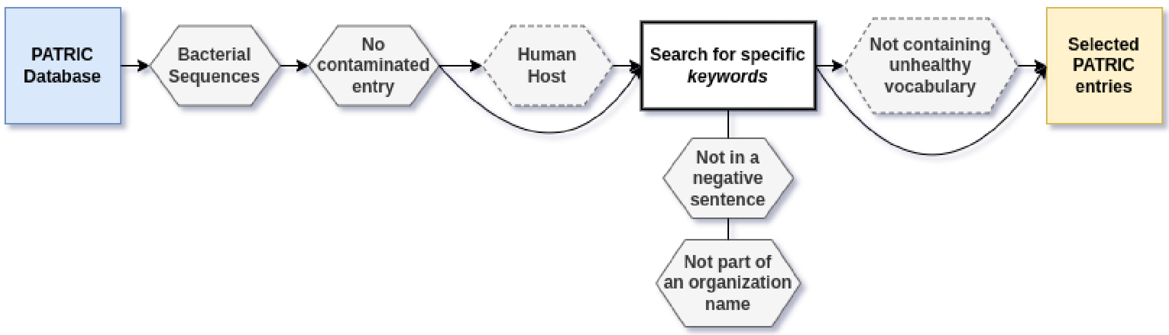


**Figure S8: Scheme of selecting entries from the PATRIC database.**

Certain steps are skipped depending on the phenotype searched. Only entries for pathogenic capacity and “Non-pathogen” and “Microbiota” are required to have “Human host”. Only entries for non-pathogens are required to “Not contain unhealthy vocabulary”.

| Virulence Function | Protein (UniRef50 ID) |
|---|---|
| Motility | **Flagellum**: A0A377CE65, UPI002021049C, A0A376J6R9, P24216, A0A2H9GEC1, A0A658Y519, A0A376MRW7, UPI00223FC8E7, A0A376Q7N2, P29744, A0A3S4IIY0, Q32DE5 |
| Adhesion to host tissue | **Adhesin**: A0A1X3JJ14, A0A2X1K134, A0A484X4U5, A0A376KQL1, Q32DV6, A0AAE9PRL0, A0A376KTK2, A0A376KRZ5, A0A376DA64. **GBS Binding surface protein**: A0A943QU00. **Anchor protein**: A0A3S4NG01. **DUF1542 domain**: UPI0028DC8763. **Histidine triad domain**: A0A428HT79. **Fimbria**: A0A8T9CK35, UPI000AECBA48, A0A377ZUH8, A0A377WDJ6, A0A0C7K737, A0A377XIZ7. **Choline binding**: J7TSG7. **VWFA:** A0AA37CTN4, UPI002AB5021D. **WxL**: A0A828QQA5, Q837W5, A0A829FD10, UPI0035A3D603, A0A829FF60, Q82ZI2, V7ZJ64. **VWA**: UPI002AB5021D. **Others**: A0A380E1C1, R2MC72, A0A4U3KFC0 |
| Colonization of host tissue | **N-acetylmuramoyl-L-alanine amidase**: A0A380EPW8, UPI002285B5FC. **Septum formation**: A0A1L7CPP2. **Lipase**: A0A2X3CBG0. **Others**: A0A380EPW8, Q8DR60 |
| Intracellular invasion | **Pullulanase**: UPI00291062F7, A0A4J1X9X5, Q8KLP1. **Internalin**: A0A930AN31. **Fibronectin**: A0A2X2YBB1, A0A9N8IGR9. **LPXTG**: A0A7T3VYA2, A0A2G0E6Y4. |
| Biofilm synthesis | **Cellulose synthase**: P37650, A0A6L3Y0A4, A0A2X3CTY4, A0A8D6Q3D5. **Formate creation**: A0A655BN26. **Glucosyl transferase:** UPI00056B38CE, A0A943QTQ6 |
| Immune system interaction | **Lipopolysaccharide**: P31554, A0A484VTJ8. **Deubiquitination**: A0A3P5DT60. **LCFA transporter**: A0A834ME79, A0AAP5MRU6. **Defense against biomolecules**: A0A4R3ZDQ6, UPI0025CEC918. **Rh**: A0A447RJP7. **Others**: UPI001E5967B5 |
| Transport/ secretion | **Autotransporter**: Q32J02, P45508, A0A8S0G0G0, A0A377EPS3, A0A376FG28, UPI0007B3DD72, UPI0009A527D6, UPI000459ED4E. **Autolysin**: P06653, A0A2S1CXQ8. **Porin**: P37681**. Secretion System (type VI)**: UPI0002C997FB, UPI002F3FDB27, R9RHK6, UPI00070728EE, F0KJ96. **Membrane permeability**: A0A447RZ74, A0A2X3EA31, UPI0030EB7CB2, UPI003523C938, UPI0021C56F87. **HlyD**: UPI002FDFCD5E. |
| Exotoxin | **Rhs toxin**: UPI00111578E0, P16917, P16919, UPI00025CA3E5, UPI0013AF20F2, G1VC94. **ShET2**: A0AAN3NUG4, C3SDH0. **Serralysin**: Q03023. |
| Stress survival | **Hydroxylamine reductase:** P75825. **D-alanyl-D-alanine carboxypeptidase**: T0U2D3. **Murein sacculus**: A0A7H4MK65. **Amine enzymes**: N9L189, A0A7H4P739 |
| Genetic material repair | **Nucleases:** A0A024L2P3, A0A377E4I2, A0A376W4B3, P15032, A0A376S820, A0A4T4CR35, A0A376PC22, A0A2X3F037, A0A377WIZ0, A0AAE4DP17, A0A4P0Y4I4. **Others**: A0A932ZA97, UPI002ED9F1EA, A0A377ZX21, A0A2X3KGH4, A0A376W4B3, A0A6L3XYV6, A0A8B3UZE9, UPI000D761D4D. |
| Pathogenic metabolism | **Nutrient acquisition:** UPI001C4F5A5B, N2A4B6, Q837L8. **Glucose catabolism**: W1DHM1, A0A377ATJ5, A0A3A1Y1R6, Q55264, U2J3Q1, A0AA42MDJ3, A0A4P0Y529, W9BGR5, |

| | |
|---|---|
| | A0A3E1ZXQ6, K1J8L7, A0A377M186, A0A519EXL9, A0A4U9HVX7, UPI0021CF0FDC, UPI002FF0DFE0. **Iron intake:** Q8XAS4, A0A2X1J1L9, A0A3L0W265, A0A2X3EIW7, A0A484ZBD7, Q9I157, Q9HV89, A0A0C7KJM8, A0A919I0T4, Q9I648, UPI0021564FC1, A0A378BUV2. **Hydrolases**: K1JCP5, A0A2X1MZ07, P25718, A0A3N6V9B5, UPI002911E5A0, UPI0024955634, S7WUG3, UPI00352BF89C, UPI000A877F0A, UPI0023E440B1, UPI00036ACD60, Q833V2, UPI0030A8EE89, A0A377YW83, UPI001CBDD196. **Anaerobic lifestyle**: P37342, P28903, A0KHM1, A0A3S4IC06, A0A2X3KGH4, A0A378A252. **Intracellular lifestyle**: A0A447S040. **Amine oxidase**: A0A385EWU8, A0A377ZUN0, A0A377D2Z8, P46883. |
| Others | **Peptidoglycan formation**: UPI0002B55144, F4NA87, A0A376GVY2. **Synthase of virulence factors**: A0A3P5DS41. **Virulence regulation**: UPI000A4D6E57. **Antimicrobial resistance**: UPI000691E8B2. **Ig-like**: Q833X5, R3KJG5, A0A3F3NQX1 |

**Table S1: Selected UniRef and Uniprot IDs.**
The highlighted proteins from genomes predicted to have pathogenic capacity in the Test-NovelSpecies and Test-2024Strains datasets were aligned against the UniRef50 database. The proteins are identified with UniProt and UniRef accession codes, as they appear in the UniRef50 database. The most frequent hits are listed in this table, along with their roles as virulence factors and in metabolic pathways.

| **HyperParameter** | **Range Parameters** | **Chosen Parameters** |
|---|---|---|
| Incorporate amount proteins | Concat, Add, False | Concat |
| Learning Rate | 5e-05, 1e-04, 5e-04, 1e-03, 5e-03 | 5e-04 |
| Sequential Dropout | 0.4, 0.3, 0.2 | 0.3 |
| Amount ConvNext Blocks | 2, 3, 4 | 2 |
| ConvNext Dimensions | 64, 128, 256, 512 | 256 |
| Stochastic Depth | 0.1, 0.2, 0.3, 0.4 | 0.3 |
| Attention Dimensions | 64, 128, 256, 512 | 64 |
| Attention Dropout | 0.3, 0.4, 0.5, 0.6 | 0.6 |

**Table S2: Hyperparameters selected through Hyperparamter optimization.**

| **Cluster Number** | **Most common Species** |
|---|---|
| 0 | *Helicobacter pylori* |
| 1 | *Campylobacter jejuni, Campylobacter coli* |
| 2 | *Providencia alcalifaciens, Proteus mirabilis* |
| 3 | *Escherichia coli, Klebsiella pneumoniae, Enterobacter hormaechei, Klebsiella variicola, Enterobacter cloacae, Citrobacter freundii, Salmonella enterica* |

| | |
|---|---|
| 4 | *Burkholderia cepacia, Burkholderia pseudomallei* |
| 5 | *Haemophilus influenzae, Haemophilus haemolyticus* |
| 6 | *Streptococcus mutans* |
| 7 | *Mycobacteroides abscessus, Mycobacterium avium* |
| 8 | *Streptococcus pneumoniae* |
| 9 | *Clostridioides difficile, Clostridium perfringens* |
| 10 | *Vibrio parahaemolyticus, Vibrio cholerae* |
| 11 | *Campylobacter concisus* |
| 12 | *Neisseria meningitidis* |
| 13 | *Leptospira interrogans, Legionella pneumophila* |
| 14 | *Stenotrophomonas maltophilia, Achromobacter xylosoxidans* |
| 15 | *Acinetobacter junii, Acinetobacter haemolyticus* |
| 16 | *Acinetobacter baumannii, Acinetobacter bereziniae* |
| 17 | *Streptococcus salivarius, Streptococcus sobrinus, Streptococcus sanguini* |
| 18 | *Clostridium sp., Clostridium perfringens* |
| 19 | *Pseudomonas aeruginosa* |
| 20 | *Pseudomonas aeruginosa, Pseudomonas putida* |
| 21 | *Aeromonas veronii, Aeromonas hydrophila, Aeromonas caviae* |
| 22 | *Streptococcus oralis, Streptococcus mitis, Streptococcus sanguinis* |
| 23 | *Streptococcus pyogenes, Streptococcus parasanguinis, Streptococcus anginosus* |
| 24 | *Staphylococcus aureus, Staphylococcus pseudintermedius* |
| 25 | *Listeria monocytogenes, Enterococcus casseliflavus* |
| 26 | *Enterococcus faecium* |
| 27 | *Gardnerella vaginalis* |
| 28 | *Enterococcus faecalis* |
| 29 | *Staphylococcus epidermidis, Staphylococcus haemolyticus* |
| 30 | *Bacteroides uniformis* |
| 31 | *Lactobacillus spp., Clostridium spp., Neisseria spp.* |

| | |
|---|---|
| 32 | *Corynebacterium spp.* |
| 33 | *Porphyromonas gingivalis, Bacteroides stercoris* |
| 34 | *Bacteroides spp.* |
| 35 | *Bacteroides fragilis* |
| **Table S3: Main bacteria species on each of the clusters shown in the Figure S4** | |
